# PAE Viewer: A webserver for the interactive visualization of the predicted aligned error for multimer structure predictions and crosslinks

**DOI:** 10.1101/2023.03.06.531253

**Authors:** Christoph Elfmann, Jörg Stülke

## Abstract

The development of AlphaFold for protein structure prediction has opened a new era in structural biology. This is even more the case for AlphaFold-Multimer for the prediction of protein complexes. The interpretation of these predictions has become more important than ever, but it is difficult for the non-specialist. While an evaluation of the prediction quality is provided for monomeric protein predictions by the AlphaFold Protein Structure Database, such a tool is missing for predicted complex structures. Here, we present the PAE Viewer webserver (http://www.subtiwiki.uni-goettingen.de/v4/paeViewerDemo), an online tool for the integrated visualization of predicted protein complex using a 3D structure display combined with an interactive representation of the Predicted Aligned Error (PAE). This metric allows an estimation of the quality of the prediction. Importantly, our webserver also allows the integration of experimental cross-linking data which helps to interpret the reliability of the structure predictions. With the PAE Viewer, the user obtains a unique tool which for the first time allows the intuitive evaluation of the PAE for protein complex structure predictions.

**GRAPHICAL ABSTRACT:** The Predicted Aligned Error (PAE) is an indication of the reliability of protein structure predictions. The PAE Viewer webserver allows the intuitive and interactive evaluation of complex structure predictions.

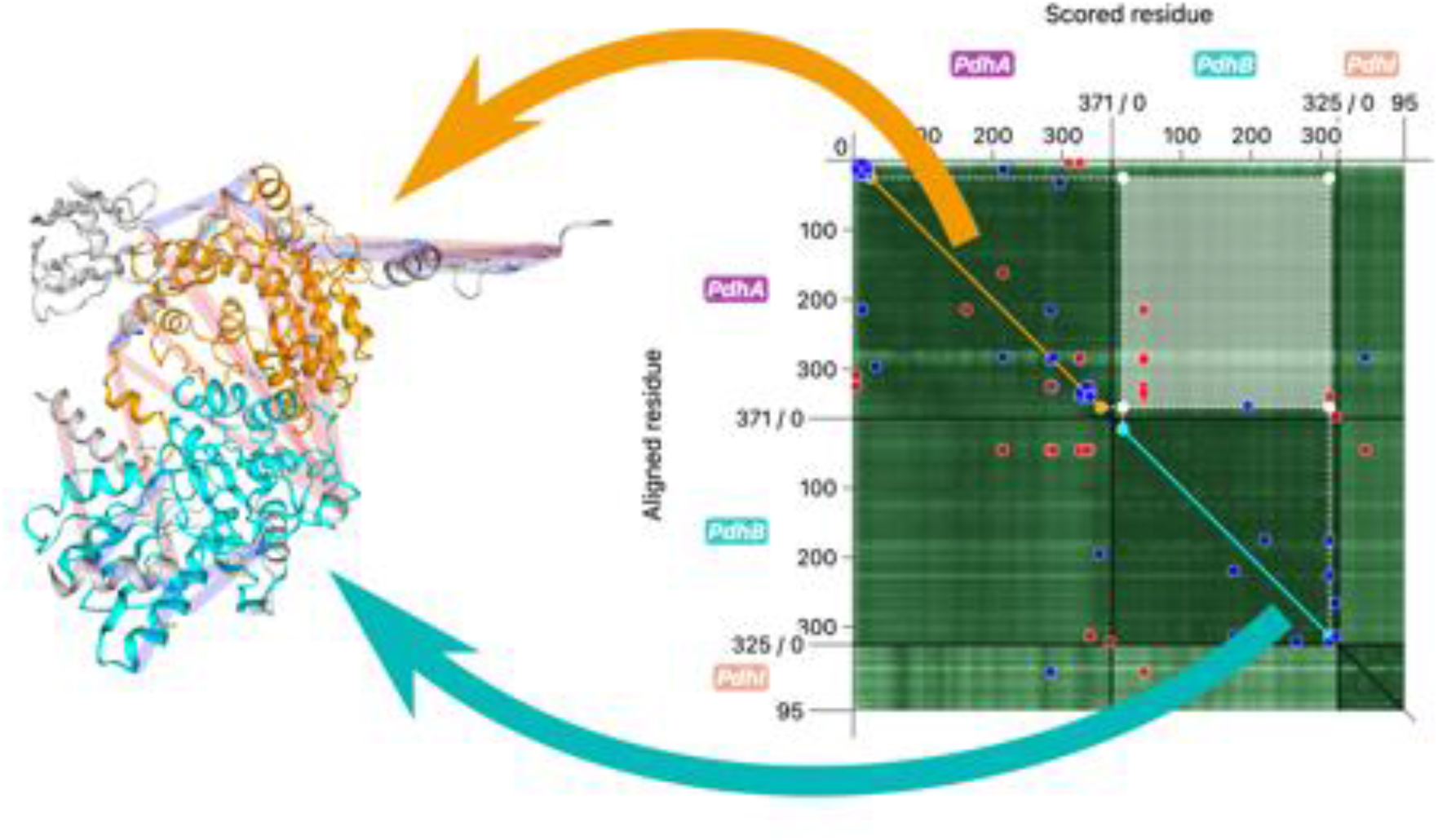

## INTRODUCTION

Many proteins act not as individual molecules but are part of larger complexes. This is particularly the case for all aspects of the replication and expression of genetic information that involve multimeric DNA polymerases, RNA polymerases and ribosomes. Also, many metabolic reactions depend on the formation of protein complexes such as ATP synthesis by a multi-subunit ATPase.

The investigation of these complexes is a major issue in current biology. Protein-protein interactions have traditionally been identified in different ways, (i) direct co-purification of unknown proteins with a specific target protein, (ii) two-hybrid assays that reconstitute enzyme activities as a result of interaction of proteins coupled to separated domains of the assay protein, and (iii) direct biochemical analysis of the interaction of candidate partners. Recently, the proteome-wide identification of protein complexes by protein cross-linking or co-fractionation coupled to mass spectrometry was introduced (1, 2). The further investigation of protein complexes has two major aspects: On one hand, the function of the interacting partners should be identified. In this case, interactions of unknown proteins with proteins of known functions paves the way to the development of hypotheses that can be experimentally addressed. In this way, the identification of interactions that involve unknown proteins is a major input to elucidate the functions of these often poorly studied proteins (3, 4). On the other hand, the analysis of the structure of the complexes and the analysis of the molecular details of the interaction is very important for a complete understanding.

Recently, the analysis of protein structures has been revolutionized by the introduction of AlphaFold, an artificial intelligence-based tool for the computational prediction of protein structures (5). With AlphaFold, structures of all proteins that can be deduced from genome databases have been predicted and have been included in the AlphaFold Protein Structure Database (AlphaFold DB) (6). However, the prediction of protein complexes is more challenging. A novel iteration of AlphaFold, AlphaFold-Multimer, has made the accurate prediction of small protein complexes feasible (7). Since its release, the prediction of larger and larger complexes has made significant progress (8). With these advances, the interpretation of the reliability of these computational predictions has become more and more urgent. For this purpose, AlphaFold and its descendants provide several metrics which allow the quantitative assessment of the quality of structure prediction. One of these indicators is the Predicted Aligned Error (PAE), which is a measure for the confidence in the relative positions and orientations of parts of the predicted structures. A 2D plot of the PAE is prominently featured on the AlphaFold DB pages, next to the three-dimensional structure view for each of the presented predictions of the individual monomeric proteins.

Structural proteomics by combining protein crosslinks and mass spectrometry provides an additional layer of information. The crosslinks indicate which residues are located in close vicinity. This can be relevant for individual proteins but also for protein complexes. Importantly, this approach even allows the detection of conformational changes of protein under different conditions. For protein complexes, the combination of AlphaFold-Multimer predictions and experimental data from structural proteomics provides an unprecedented view of the molecular organization of the complexes. Indeed, in a recent study on the *Bacillus subtilis* interactome, a so far unknown protein that interacts with two subunits of pyruvate dehydrogenase was identified. This protein, then renamed PdhI, was found to inhibit the enzyme activity of the pyruvate dehydrogenase. Structural modeling suggested that the protein interferes with catalysis by protruding into the active center of the enzyme. Site-directed mutagenesis based on this hypothesis finally confirmed this proposed mechanism of inhibition of the pyruvate dehydrogenase (2).

The experimental identification of crosslinks between specific regions of interacting proteins can provide independent validation of AlphaFold-Multimer complex predictions. For individual proteins, this experimental evidence can give important hints about the validity of the predicted conformation. The same is true for complexes, where crosslinks between subunits additionally provide information on the shared interface. Thus, the integration of this experimental crosslink information can help to assess the accuracy of these predictions.

To help researchers with the interpretation of complex predictions, we developed the PAE Viewer webserver. With this online tool, users can view the 3D representation of a predicted protein complex, corresponding amino acid sequences as well as an interactive display of the PAE. All of these components work together, so user interactions with one of them are reflected by the others. While this functionality is similar to the one of AlphaFold DB entry pages, particular focus was put on the representation and interactivity of the PAE display, the *PAE Viewer.* While the PAE plot on AlphaFold DB is limited to protein monomers, our implementation can display the PAE of both monomers and protein complexes in a comprehensive manner. Additionally, we equipped it with advanced controls to facilitate an intuitive exploration of the PAE. By incorporating crosslinking data into the display, the PAE Viewer also allows for the integration of experimental data. With this unique combination of features, our webserver provides a comprehensive tool to interpret the quality of multimeric structure predictions. A version of the PAE Viewer is currently integrated into the database *SubtiWiki* on the model organism *B. subtilis* (9), where a predefined selection of predicted protein complex structures is presented. In contrast, the PAE viewer webserver can be used for uploading any predicted custom structure that is of interest to the user.

## IMPLEMENTATION

### The Predicted Aligned Error

AlphaFold, in addition to the predicted structure, provides several metrics which allow to better assess the prediction quality. One of these is the Predicted Aligned Error (PAE), a measure for the confidence in the relative position of two residues within the predicted structure (6). The PAE for a pair of residues *x* and *y* is defined as the expected positional error in Ångströms at *x* if the predicted and actual structures are aligned at *y*. This can provide valuable information about the reliability of relative position and orientations of different domains: if the PAE between the contained residues is high, AlphaFold predicts these domains to be accurately oriented; in turn, a low value indicates limited reliability of the predicted domain orientation. Correspondingly, the same is true for AlphaFold-Multimer predictions with regard to different chains of the model. In this context, a high PAE between the residues of different chains indicates a reliable prediction of the shared interface (7).

### Overview of the webserver functionality

When assessing the quality of structure predictions, it can be difficult to interpret the PAE and its relation to the predicted structure. For this reason, many applications concerned with the evaluation of these predictions feature 2D plots to illustrate the PAE. In case of the AlphaFold DB entry pages, these displays even provide functionality to select parts of the structure and the associated PAE values. However, as mentioned before, the implementation by AlphaFold DB has certain shortcomings, such as the limitation to monomer predictions, among others. For this reason, we developed the PAE Viewer webserver, which is suitable for the evaluation of protein multimer predictions and which features an advanced interactive display of the PAE. The website allows users to upload and view AlphaFold-Multimer predictions and the associated PAE in an intuitive way.

The PAE Viewer webserver page presents an integrated view of an AlphaFold-Multimer structure prediction (Fig. 1). In addition to the PAE Viewer itself, the page features interactive displays of the associated amino acid sequences as well as the 3D structure. The sequence viewer displays the amino acid sequences of the chains contained in the predicted multimer. By using the mouse, the user can select individual amino acids or continuous ranges, which in turn are highlighted in the 3D structure viewer. The latter is based on *NGL viewer,* a web-based tool for molecular visualization (10). The structure viewer shows a 3D representation of the predicted multimer, which the user can interact with by rotating, dragging or zooming with the mouse. It also features information on additional prediction quality metrics and offers options for the choice of color schemes and display of crosslinks. The PAE Viewer, the sequence viewer and the 3D structure viewer work in conjunction with each other, so user interactions with one of the viewers are reflected by the other displays.

**Figure 1.**
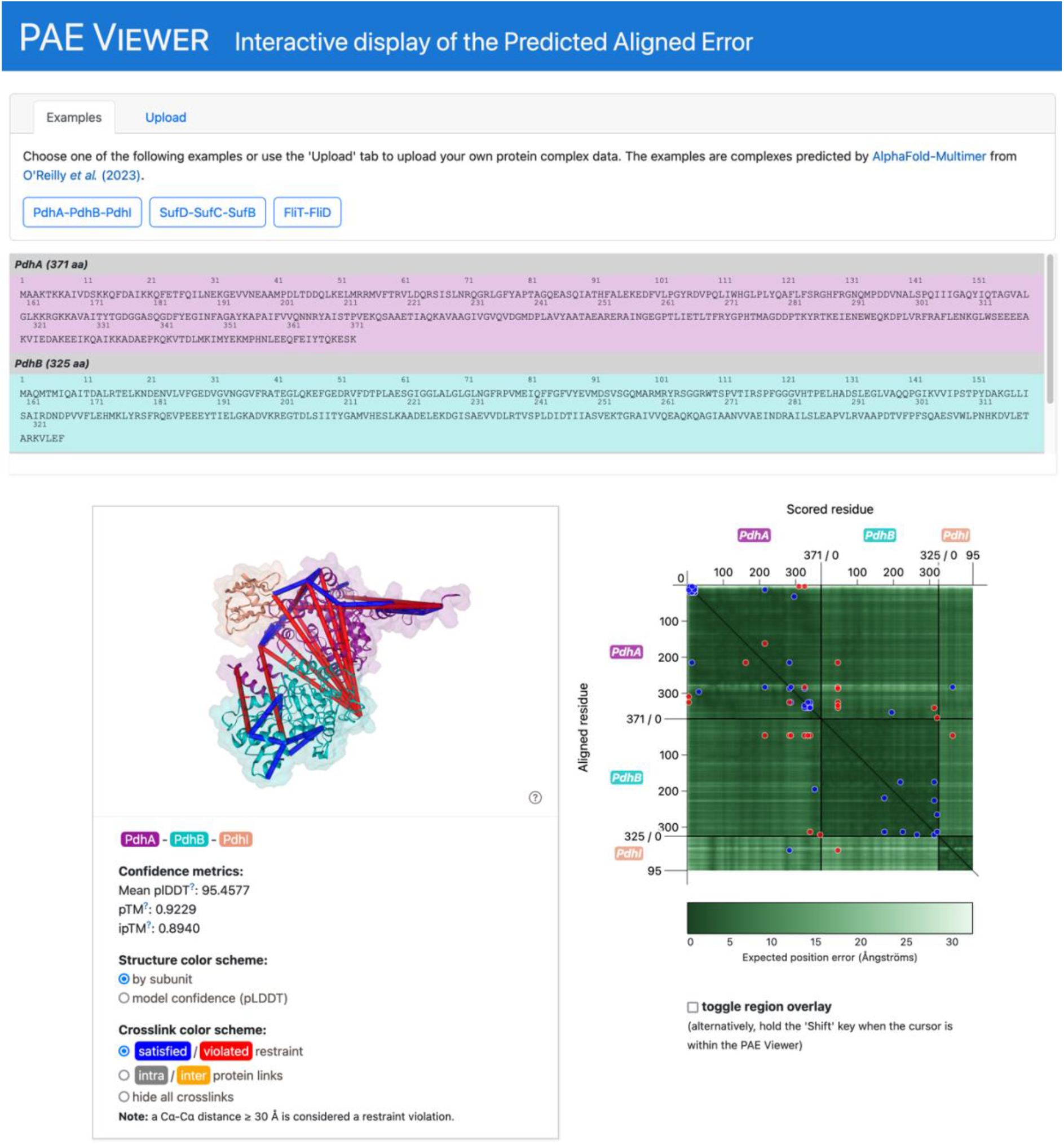
Overview of the PAE Viewer webserver page. The panel at the top of page allows to select structure data from several examples and to upload custom data. Below, the dynamic sequence viewer presents the amino acid sequences of the viewed multimer prediction. Further down on the left, a 3D structure viewer shows a molecular representation of the predicted multimer. Additionally, quality metrics and presentation options are provided. On the right, the PAE Viewer is embedded.

At the top of the page, the user can choose from a selection of example structures to test the page functionality. Furthermore, an upload form allows users to provide their own structural data. The featured example data includes structure predictions, crosslinking data and quality metrics. They are derived from a recent global interaction study on *B. subtilis,* where AlphaFold-Multimer was used in combination with experimental crosslinking and co-fractionation analyses (2). This approach resulted in the prediction of high-quality models of numerous potential protein-protein interactions, of which three were chosen as example data. The functionality of the webserver page is entirely provided by the client-side web application, so no exchange of information with our server is performed when users “upload” data.

### Representation of data with the PAE Viewer

The PAE Viewer was designed to facilitate the interpretation of the PAE for multimeric predictions. To this end, a two-dimensional, interactive plot was developed which features intuitive selection of structure parts, as well as integration of crosslinking data. Fig. 2A shows the PAE Viewer for one of the example complexes, the PdhA-PdhB-PdhI trimer in *B. subtilis.*

**Figure 2.**
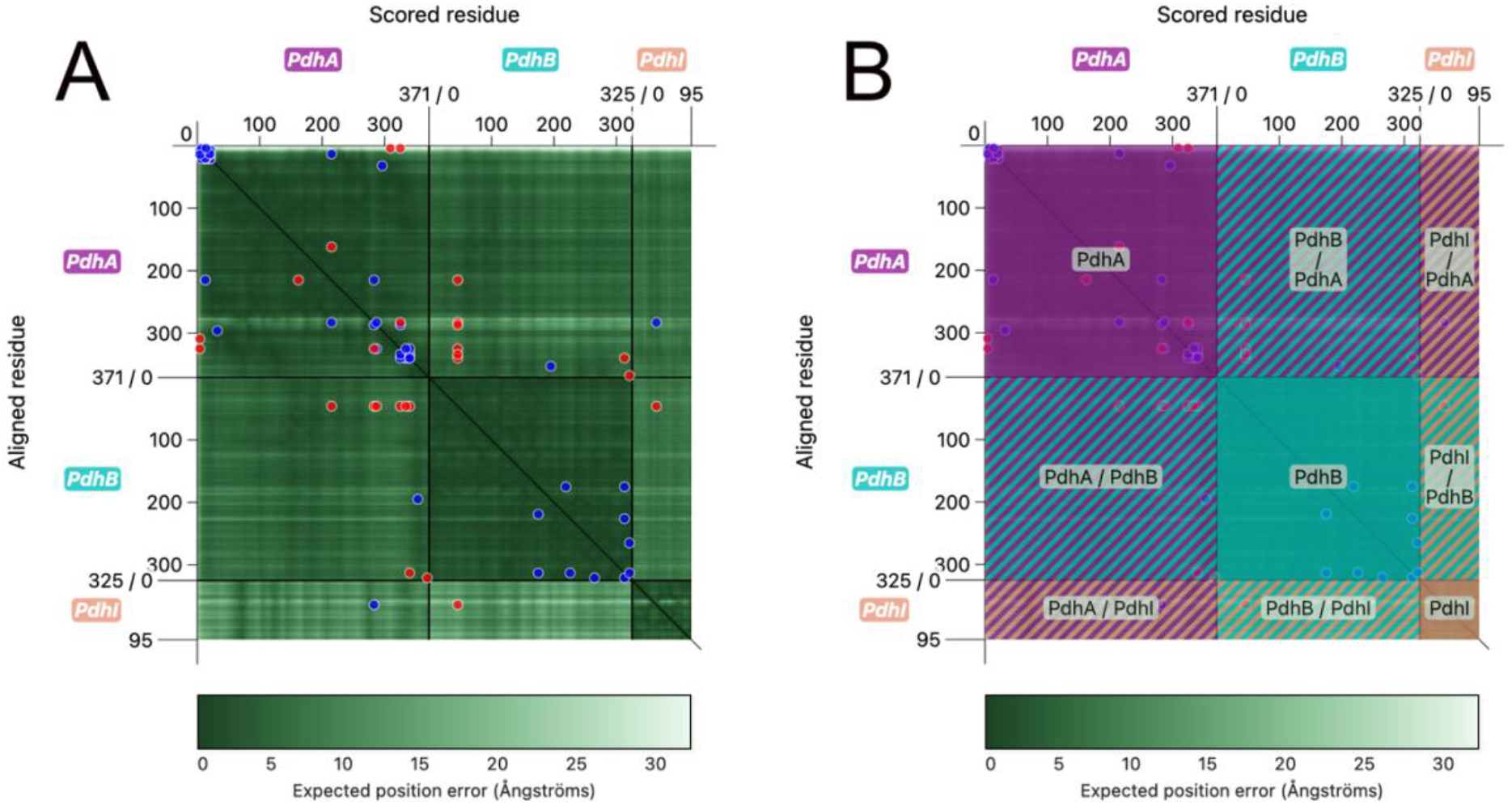
The PAE Viewer display. A: The interactive PAE Viewer plot, which features a PAE heatmap and additional graphical elements. B: The PAE Viewer with enabled “region overlay”, which allows to highlight the plot segments corresponding to different chains or interfaces.

As can be seen, the representation of the PAE is similar to the display used by the AlphaFold DB. However, additional graphical elements were added to distinguish between the different chains of the multimer, which are not present in AlphaFold DB, as the latter currently only features monomer predictions. Furthermore, our PAE Viewer allows the integration of crosslinking data, where circular markers indicate crosslinks between residues.

The 2D plot displays the PAE at the scored residue *x* with regards to the aligned residue *y* as a so-called heatmap, where the value of the PAE is indicated by the color. *A* dark green corresponds to a low PAE, which indicates a high reliability of the relative position of the residues. Conversely, a lighter color corresponds to a lower confidence. This color scheme is commonly used for the PAE, as for instance by AlphaFold DB. The expected positional error corresponding to the PAE is capped at 31.75 Å, as defined by the AlphaFold-Multimer output.

The x and y axes are segmented by the multimer chains, so the position of a residue within a subunit can be easily seen. Special axis ticks also indicate the total sequence length of the individual chains. Additional color-coded labels denote the names of the chains. The color scheme is consistent for the PAE viewer, the sequence viewer and the 3D models. A colorblind-friendly scheme based on the Okabe/Ito palette was used (https://web.archive.org/web/20230223220710/https://jfly.uni-koeln.de/color/).

The heatmap itself is segmented into a grid, where rectangular parts of the plot correspond to different chains, or interfaces between two chains. These can be highlighted and labeled by toggling the *“region overlay”,* which applies a color-coded mask to the plot as shown in Fig. 2B.

In addition to the PAE itself, crosslinking data can be integrated. A crosslink between two residues at *x* and *y* is displayed as a pair of circular markers at (*x*, *y*) and (*y*, *x*) within the plot. Color coding is used to denote satisfaction (blue) or violation (red) of crosslink restraints. In the example (Fig. 2A), a distance restraint was imposed, where a Cα-Cα distance ≥ 30 Å was considered a restraint violation, as such a distance was deemed physically impossible for the used crosslinker. Combined with the PAE, the existence of crosslinks with either satisfied or violated distance restraints provides additional hints about the confidence of the conformation and orientation of structure parts.

### Interactivity of the PAE Viewer

Further differences to the PAE plot of the AlphaFold DB arise when comparing interactivity of selections. Fig. 3 shows the state of the webserver page when a part of the heatmap is selected.

**Figure 3.**
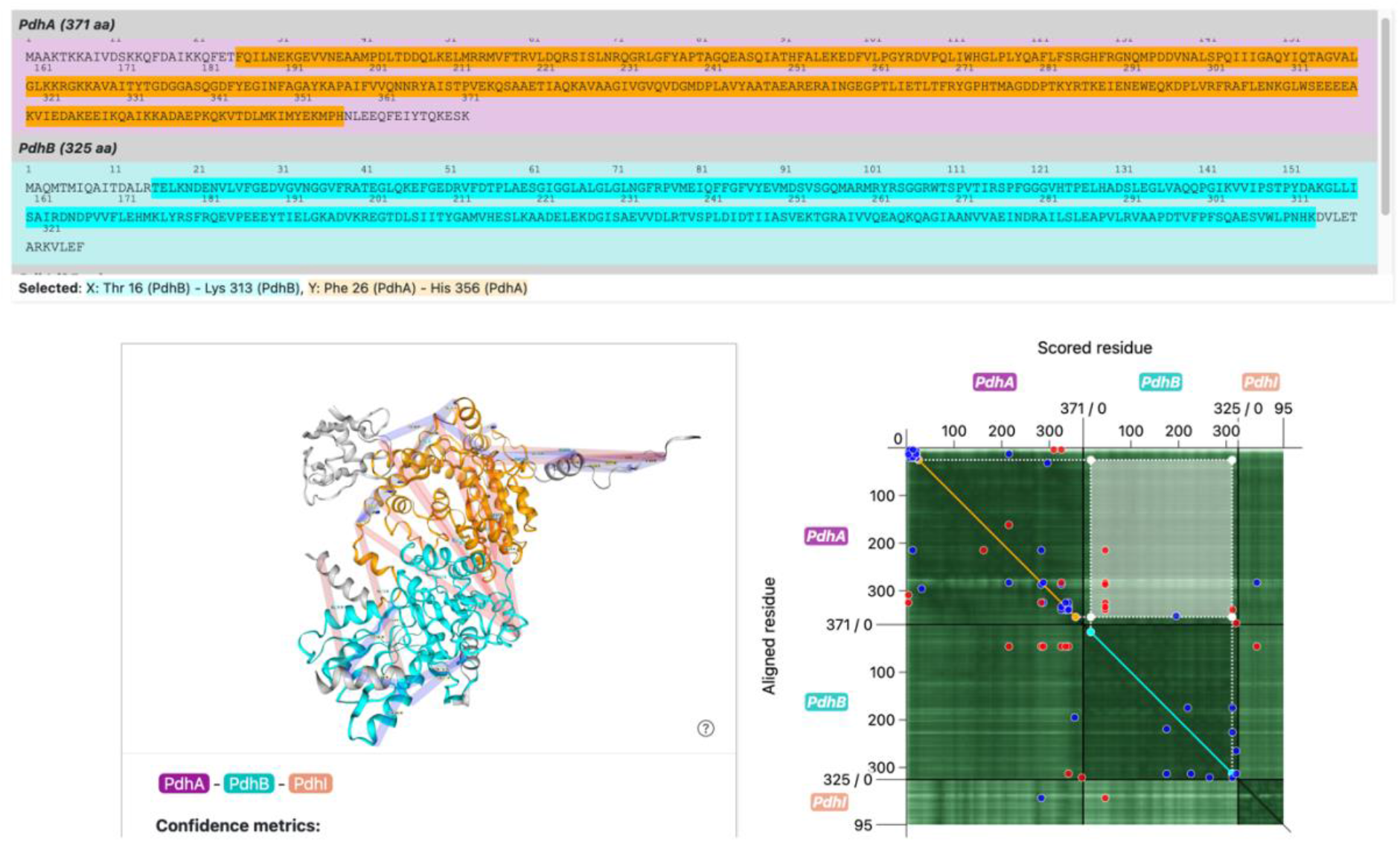
Selection functionality of the PAE Viewer. When a selection is performed on the PAE Viewer, the corresponding parts of the multimer are highlighted in the sequence viewer as well as the 3D structure viewer using a consistent color scheme.

The selection functionality of the PAE viewer was designed to explore the PAE and the corresponding parts of the multimer prediction intuitively. When clicking and holding the left mouse button while dragging the cursor across the heatmap, a rectangular area of the plot can be selected as demonstrated in Fig. 3. Such a selection corresponds to the PAE of one range of residues with regards to another. Both ranges are projected onto the diagonal of the plot to illustrate the relation of the selected ranges within the sequence. The *x* range, which corresponds to the selection of scored residues, is marked cyan, while the *y* range of aligned residues is marked orange. The same color scheme is applied to the sequence viewer and the 3D structure viewer, which makes it easy for the user to identify the different parts of the selection. By looking at the color-coded PAE values in the heatmap, the user can get an idea about the reliability of the orientation of the cyan-colored part of the model with regards to the orange-colored one. Additionally, the user can see the corresponding amino acid sequences at one glance.

On AlphaFold DB, selections with the featured PAE plot also allow highlighting of different sequence and 3D structure parts. However, the AlphaFold DB display highlights the ranges of scored and aligned residues the same way, and also includes residues *between* the two selected ranges. This makes the identification of the corresponding parts in the 3D structure viewer impossible. In contrast, the PAE Viewer makes a distinction between the ranges which are highlighted in different colors. This makes it easier to identify the relation of the highlighted structure parts and sequences with the selected PAE.

Aside from the selection of rectangular areas of the PAE heatmap, the PAE Viewer allows different selections to be made. Crosslinks, which are denoted by circular markers, can be clicked to highlight corresponding representations in the 3D structure viewer. The togglable region overlay also features interactivity. When clicking an area of the overlay, the corresponding chain or interface is highlighted in the 3D structure viewer as well as the sequence viewer, using the multimer color scheme.

## CONCLUSION

Protein complexes play an essential role in cellular life, and the investigation of their functions is a major objective in biology. Structural proteomics can provide crucial insights into the workings of these molecular machineries, but experimental approaches to their determination present a demanding challenge. However, with the rise of powerful machine-learning algorithms such as AlphaFold and its offspring tools, the accurate structure prediction of larger and larger protein complexes has become a reality. In turn, the interpretation of these computational predictions and the evaluation of their quality has become more important than ever. While programs such as AlphaFold can provide quantitative measures for the confidence of their predictions, the interpretation can be difficult for the non-specialist. One of these metrics, the Predicted Aligned Error (PAE), is an important indicator for the accuracy of orientation of parts of the structures to one another. In addition to quality estimates generated by AlphaFold, experimental validation is a vital step to verify the reliability of complex structure predictions. Approaches such as crosslinking analysis can deliver important hints about the molecular organization of protein complexes.

The PAE Viewer webserver provides an intuitive tool to interactively explore the quality of AlphaFold-Multimer predictions by combining the PAE with sequence, structure, and crosslink information. The PAE Viewer display itself offers unique features, such as the support of multimeric structure predictions, the integration of crosslinking data, and advanced controls. The visualization and interactive behavior were designed to enable intuitive usage.

## AUTHOR CONTRIBUTIONS

Christoph Elfmann: Conceptualization, Implementation, Writing—original draft. Jörg Stülke: Funding acquisition, Writing—review & editing.

## ACKNOWLEDGEMENTS

We are grateful to Andrea Graziadei, Francis O’Reilly and Juri Rappsilber for helpful discussion.

## REFERENCES

1. O’Reilly, F.J., Xue, L., Graziadei, A., Sinn, L., Lenz, S., Tegunov, D., Blötz, C., Singh, N., Hagen, W.J.H., Cramer, P., et al. (2020) In-cell architecture of an actively transcribing-translating expressome. Science 369, 554–557. https://pubmed.ncbi.nlm.nih.gov/32732422/ https://doi.org/10.1126/science.abb3758 https://www.ncbi.nlm.nih.gov/pmc/articles/PMC7115962/

2. O’Reilly, F. J., Graziadei, A., Forbrig, C., Bremenkamp, R., Charles, C., Lenz, S., Elfmann, C., Fischer, L., Stülke, J., & Rappsilber, J. (2023) Protein complexes in cells by AI-assisted structural proteomics. Mol. Syst. Biol., 19, e11544. https://pubmed.ncbi.nlm.nih.gov/36815589/ https://doi.org/10.15252/msb.202311544

3. Kustatscher, G., Collins, T., Gingras, A.C., Guo, T., Hermjakob, H., Ideker, T., Lilley, K.S., Lundberg, E., Marcotte, E.M., Ralser, M. and Rappsilber, J. (2022) Understudied Proteins: opportunities and challenges for functional proteomics. Nat. Methods 19, 774–779. https://pubmed.ncbi.nlm.nih.gov/35534633/ https://doi.org/10.1038/s41592-022-01454-x

4. Wicke, D., Meißner, J., Warneke, R., Elfmann, C. and Stülke, J. (2023) Understudied proteins and understudied functions in the model bacterium *Bacillus subtilis* – a major challenge in current research. Mol. Microbiol. In press. https://doi.org/10.22541/au.167534123.32069234/v1

5. Jumper, J., Evans, R., Pritzel, A., Green, T., Figurnov, M., Ronneberger, O., Tunyasuvunakool, K., Bates, R., Žídek, A., Potapenko, A., et al. (2021) Highly accurate protein structure prediction with AlphaFold. Nature, 596, 583–589. https://pubmed.ncbi.nlm.nih.gov/34265844/ https://doi.org/10.1038/s41586-021-03819-2 https://www.ncbi.nlm.nih.gov/pmc/articles/PMC8371605/

6. Varadi, M., Anyago, S., Deshpande, M., Nair, S., Natassia, C., Yordanova, G., Yuan, D., Stroe, O., Wood, G., Laydon, A., et al. (2022) AlphaFold Protein Structure Database: massively expanding the structural coverage of protein-sequence space with high-accuracy models. Nucleic Acids Res., 50, D439–D444. https://pubmed.ncbi.nlm.nih.gov/34791371/ https://doi.org/10.1093/nar/gkab1061 https://www.ncbi.nlm.nih.gov/pmc/articles/PMC8728224/

7. Evans, R., O’Neill, M., Pritzel, A., Antropova, N., Senior, A., Green, T., Žídek, A., Bates, R., Blackwell, S., Yim, J., et al. (2021) Protein complex prediction with AlphaFold-Multimer. BioRxiv. https://doi.org/10.1101/2021.10.04.463034.

8. Bryant, P., Pozzati, G., Zhu, W., Shenoy, A., Kundrotas, P. and Elofsson, A. (2022) Predicting the structure of large protein complexes using AlphaFold and Monte Carlo tree search. Nat. Commun., 13, 6028. https://pubmed.ncbi.nlm.nih.gov/36224222/ https://doi.org/10.1038/s41467-022-33729-4 https://www.ncbi.nlm.nih.gov/pmc/articles/PMC9556563/

9. Pedreira, T., Elfmann, C. and Stülke, J. (2022) The current state of *Subti*Wiki, the database for the model organism *Bacillus subtilis*. Nucleic Acids Res., 50, D875–D882. https://pubmed.ncbi.nlm.nih.gov/34664671/ https://doi.org/10.1093/nar/gkab943 https://www.ncbi.nlm.nih.gov/pmc/articles/PMC8728116/

10. Rose, A. S., Bradley, A. R., Valasatava, Y., Duarte, J. M., Prlic, A. and Rose, P. W. (2018) NGL viewer: web-based molecular graphics for large complexes. Bioinformatics 34, 3755–3758. https://pubmed.ncbi.nlm.nih.gov/29850778/ https://doi.org/10.1093/bioinformatics/bty419 https://www.ncbi.nlm.nih.gov/pmc/articles/PMC6198858/

